# Requirement for an Otopetrin-Like protein for acid taste in *Drosophila*

**DOI:** 10.1101/2021.06.18.449071

**Authors:** Anindya Ganguly, Avinash Chandel, Heather Turner, Shan Wang, Emily R. Liman, Craig Montell

**Affiliations:** Neuroscience Research Institute and Department of Molecular, Cellular and Developmental Biology, University of California, Santa Barbara, CA 93106; Department of Biological Sciences, Section of Neurobiology, University of Southern California, Los Angeles, CA 90089

**Keywords:** sour, acid taste, Otopetrin, *Drosophila*, proton sensor, gustation

## Abstract

Many of the *Drosophila* receptors required for bitter, sugar and other tastes have been identified. However, the receptor required for the taste of acid has been elusive. In *Drosophila*, the major families of taste receptors, such as “Gustatory Receptors” and “Ionotropic Receptors” are unrelated to taste receptors in mammals. Previous work indicated that members of these major families do not appear to be broadly required acid sensors. Here, to identify the enigmatic acid taste receptor, we interrogated three genes encoding proteins distantly related the mammalian Otopertrin1 proton channel. We found that RNAi knockdown or mutation of *Otopetrin-Like A* (*OtopLA*) by CRISPR/Cas9, severely impairs the behavioral rejection of sugary foods laced with HCl or carboxylic acids. Mutation of *OtopLA* also greatly reduces acid-induced action potentials. We identified an isoform of *OtopLA* that was expressed in the proboscis and was sufficient to restore acid sensitivity to *OtopLA* mutant flies. *OtopLA* functioned in acid taste in a subset of bitter-activated gustatory receptor neurons that senses protons. This work highlights an unusual functional conservation of a receptor required for a taste modality in flies and mammals.

## Introduction

The sense of taste in some mammals is quite different from humans. Cats are strictly carnivores and are not endowed with sweet taste (1). Due to environmental pressures, the bottlenose dolphin represents an extreme example of taste deviation in mammals since they detect salt, but not sweet, bitter or other chemicals in food (2). In contrast, the fruit fly, *Drosophila melanogaster*, responds to a similar repertoire of tastes as humans despite the distant evolutionary relatedness and the enormous differences in the anatomy of their taste organs. Similar to humans, the fruit fly detects sweet, bitter, salt, amino acid, fatty acid and sour (low pH) (3). Many of the receptors involved in *Drosophila* taste have been defined (3, 4). Those that contribute to sweet and bitter sensations have been characterized most extensively (3, 4). A major theme that has emerged is that the fly taste receptors, such as the large families of “Gustatory Receptors” (GRs) and “Ionotropic Receptors” (IRs) are distinct from those that function in mammalian taste (4). Therefore, the abilities of insects and humans to respond to similar repertoires of chemicals such as sweet and bitter tastants may have emerged independently.

In mice the taste of acids depends on a proton selective channel, Otopetrin1 (Otop1), which is expressed in type III taste receptor cells (5-7). However, in *Drosophila* the receptor that is required for tasting protons has not been defined. Although low levels of some organic acids are attractive to flies and promote feeding, acids at modest and especially at high concentrations repress food consumption (8, 9). This rejection contributes to survival as it discourages the animals from eating very acidic foods in the environment that can decrease lifespan.

Currently, no broadly required acid sensor has been identified in flies. IR7a is the only known receptor that is needed to suppress feeding of an acidic compound (9). However, IR7a is very narrowly tuned as it impairs the rejection of foods with acetic acid but not HCl or any other carboxylic acid tested. Thus, IR7a cannot be a proton sensor. This receptor acts in a subset of gustatory receptor neurons (GRNs) called B GRNs that are also activated by bitter chemicals and certain other aversive compounds (3, 9). Two other IRs (IR25a and IR76b) function in GRNs in the legs for sensing acids and this ability contributes to the selection of egg laying sites (10). However, neither of these IRs appear to function in feeding decisions based on acidity (9).

In this work we set out to identify the proton sensor that is broadly required for acid taste in *Drosophila*. In contrast to other taste modalities in flies, we found that the taste of protons depends on a receptor that has a common evolutionarily origin with mice. We found that a distantly related *Drosophila* Otopetrin-Like protein (OtopLA) is required for the gustatory rejection of HCl and carboxylic acids, and promotes production of proton-induced action potentials in acid-activated B GRNs. Thus, in contrast to other gustatory modalities, acid taste is mediated through a sensor that is conserved in flies and mammals, despite the ∼800 million years that have elapsed since their common ancestor (http://www.timetree.org/) (11).

## Results

### Silencing *OtopLA* reduces gustatory repulsion to acids

To characterize the impact of HCl and organic acids on the appeal of 100 mM sucrose we conducted proboscis extension response (PER) assays. We starved control flies for a long period (24-26 hours) so that they were highly motived to feed. We then touched the major taste organ (the labellum) with sucrose and determined whether the animals extended their proboscis thereby indicating their interest in feeding. A full extension is scored as 1.0, a partial extension as 0.5 and no extension as 0. When we offered control flies sucrose only (pH 6.8), virtually all the animals fully extended their proboscis (Fig. 1*A*).

**Fig. 1.**
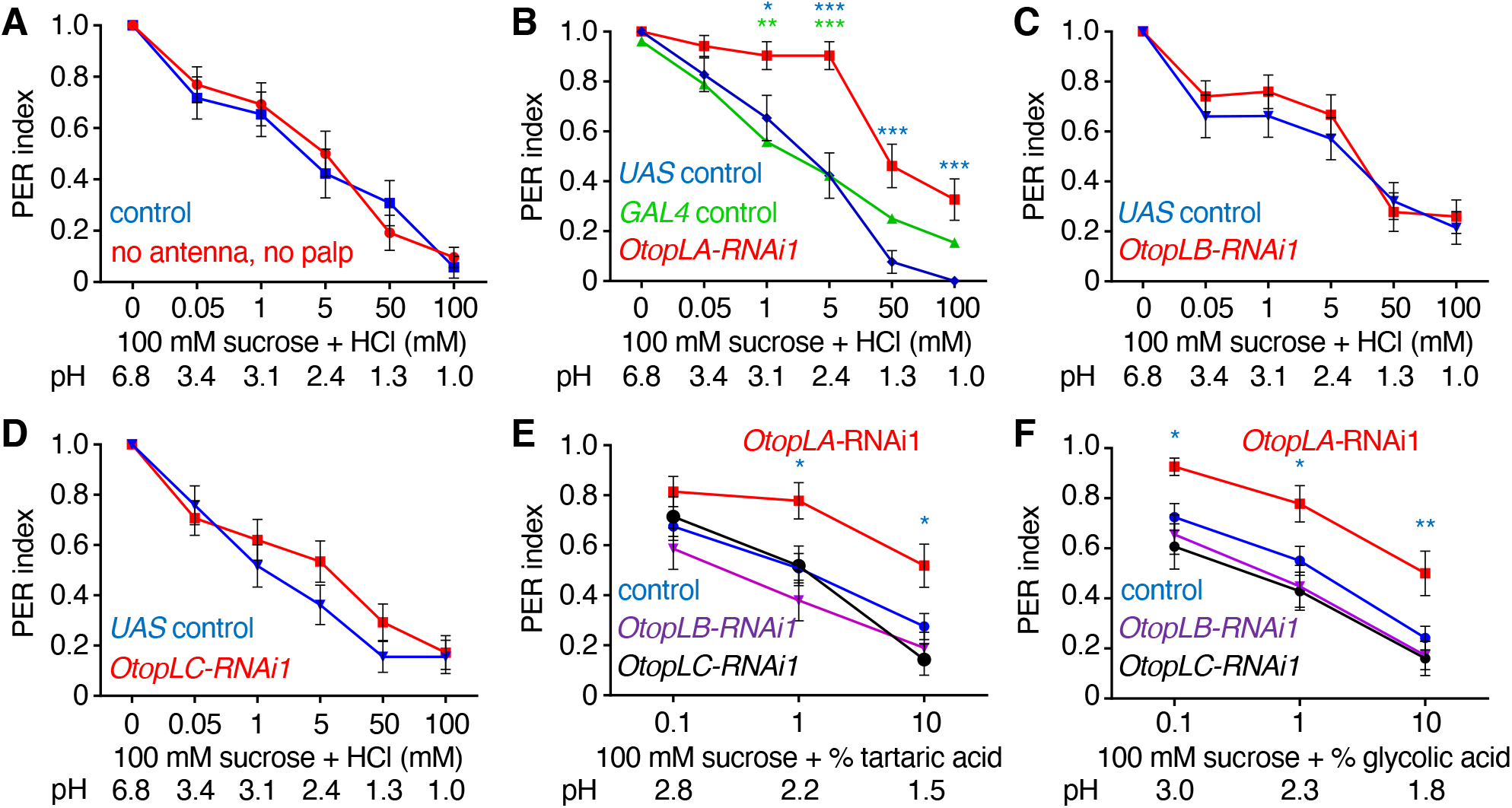
RNAi screen for *OtopL* genes required for aversion of acid taste. Proboscis extension response (PER) assays were performed by touching the proboscis with either 100 mM sucrose alone, or sucrose and the indicated concentrations of HCl (mM) or organic acids (%). The pH values are indicated in each panel. A full extension is scored as 1.0, a partial extension as 0.5 and no extension as 0. All RNAi lines (*B—F*) were generated by crossing the indicated *UAS*-*RNAi* lines to *elav-GAL4;UAS-Dcr2* flies. (*A*) Effects of removing olfactory organs on the PER responses to the indicated concentrations of HCl. Control (*w*^*1118*^) and *w*^*1118*^ flies in which the antenna and maxillary palp were surgically removed (no antenna, no palp). n=26. (*B—F*) Effects of RNAi knockdown of different *OtopL* genes on the PER responses to the indicated acids. (*B*) Effect of knockdown of *OtopLA* on the PER to sucrose plus HCl. *OtopLA-RNAi1* is *UAS-OtopLA-RNAi1* (v104973) crossed to *elav-GAL4;UAS-Dcr2* flies (n=30). The *UAS* control (n=26) and *GAL4* control (n=25) are generating by crossing *w*^*1118*^ to either *UAS-OtopLA-RNAi1* or *elav-GAL4;UAS-Dcr2*, respectively. The blue and green asterisks indicate statistically significance differences between *OtopLA* silenced flies and the *UAS* and *GAL4* controls, respectively. (*C*) Effect of knockdown of *OtopLB* on the PER to sucrose plus HCl. *OtopLB-RNAi1* is *UAS-OtopLB-RNAi1* (v101936) crossed to *elav-GAL4;UAS-Dcr2* flies (n=27). *UAS* control (n=28). (*D*) Effect of knockdown of *OtopLC* on the PER to sucrose plus HCl. *OtopLC-RNAi1* is *UAS-OtopLC RNAi* (v108591) crossed to *elav-GAL4;UAS-Dcr2* flies (n=29). *UAS* control (n=29). (*E*) Effect of knockdown of *OtopL* genes on PERs using the indicated concentrations of tartaric acid. *OtopLA-RNAi1* (n=27), *OtopLB-RNAi1* (n=29), *OtopLC-RNAi1* (n=28). The “control” is *elav-GAL4;UAS-Dcr2* flies (n= 65). The blue asterisks indicate significance differences with the control. (*F*) Effect of knockdown of *OtopL* genes on PERs using the indicated concentrations of glycolic acid. *OtopLA-RNAi1* (n=27), *OtopLB-RNAi1* (n=29), *OtopLC-RNAi1* (n=28). The “control” are *elav-GAL4;UAS-Dcr2* flies (n= 65). The blue asterisks indicate significance differences with the control. Mann-Whitney U tests. Error bars, s.e.m.s. *p < 0.05, **p < 0.01, and ***p < 0.001.

We then laced the sucrose with different concentrations of HCl thereby decreasing the pH from 6.8 to between 3.4 (50 μM HCl) and 1.0 (100 mM HCl). Control flies reduced their propensity to extend their proboscis to sucrose in proportion to increasing concentrations of HCl, as well as organic acids such as tartaric acid (Fig. 1*A* and Fig. S1*A*). Flies sense acids through both taste and smell (3). To assess a potential contribution of smell to the suppression of the PER by HCl we surgically ablated the two olfactory organs: the antenna and maxillary palp. After removing these organs, HCl and tartaric acid still diminished the PER to the same extend as intact control flies (Fig. 1*A* and Fig. S1*A*). Thus, olfaction was not needed for the flies to elicit a distaste for acids.

*Drosophila* encodes three Otopetrin-Like proteins (*OtopLA*—*C*), which share a low level of amino acid homology (20.1 to 30.6% identities) to the mouse and human Otop1 proteins (7, 12). Despite this modest sequence homology, we used RNAi to investigate if any fly OtopL was required for gustatory aversion towards acids. We used two RNAi lines for each gene and drove expression with a pan-neuronal driver (*elav-GAL4*). Both *OtopLA* RNAi lines (*OtopLA-RNAi1* and *OtopLA-RNAi2*) caused large decreases in repulsion to low pH (Fig. 1*B* and Fig. S1*B*). Between 50 μM and 5 mM HCl (pH 3.4 to 2.4) the effects of *OtopLA-RNAi* were profound as there were only minimal reductions in sucrose attraction (Fig. 1*B* and Fig. S1*B*). Even when we added 100 mM HCl (pH 1.0) to the sugar, there was significantly less acid-induced suppression of the PER relative to controls (Fig. 1*B* and S1*B*). The remaining rejection of sucrose at very low pH of 1.3 and 1.0 could possibly be due to the nociceptive response to extremely high acidity (13). In contrast to the effects of silencing *OtopLA*, RNAi knockdown of either *OtopLB* or *OtopLC* had no effect on the HCl-induced suppression of the PER to sucrose (Fig. 1 *C* and *D* and Fig. S1 *C* and *D*).

To investigate if *OtopLA* is also required for sensing carboxylic acids we added tartaric acid and glycolic acid to the sucrose and performed PER assays. The suppression by these carboxylic acids that occurred in control flies was also significantly reduced by silencing *OtopLA* at all concentrations tested (Fig. 1 *E* and *F* and Fig. S1 *E* and *F*). However, knockdown of *OtopLB* or *OtopLC* had no impact (Fig. 1 *E* and *F* and Fig. S1 *E* and *F*).

### A proboscis *OtopLA* isoform functions in acid sensation

The *OtopLA* locus is predicted to be expressed as six mRNA isoforms, which encode 916-991 amino acid proteins with 12 transmembrane segments similar to other Otop proteins (12, 14). Five isoforms share a common translation start site (*OtopLAc*— *OtopLAg*) and one begins at an alternative site in an exon unique to that isoform (*OtopLAa*; Fig. S2*A*). Using CRISPR/Cas9 we inserted the *LexA* and *mini-white* transgenes at the translation start site for *OtopLAc*—*OtopLAg* to create the *OtopLA*^*1*^ allele (Fig. S2*A*). The knock-in deleted 40 base pairs that removed residues 1-14 and shifted the reading frame (Fig. S2*A*). The deletion also disrupted expression of the mRNA (Fig. S2*B*). We introduced a similar insertion at the start site of *OtopLAa*, which removed the region encoding amino acids 1-18 and introduced a frame shift (Fig. S2*A*; *OtopLA*^*2*^).

We performed PER assays with 100 mM sucrose mixed with HCl over a range of concentrations. The repulsion to HCl was virtually eliminated in *OtopLA*^*1*^ flies at levels up to 5 mM, and there was a significant reduction in the PERs at higher concentrations (Fig. 2*A*). In contrast, *OtopLA*^*2*^ exhibited a normal rejection of HCl (Fig. S2*C*). Consistent with this finding, the *OtopLA-RNAi1*, which did not target *OtopLAa*, still disrupted rejection of HCl (Fig. 1*B*).

**Fig. 2.**
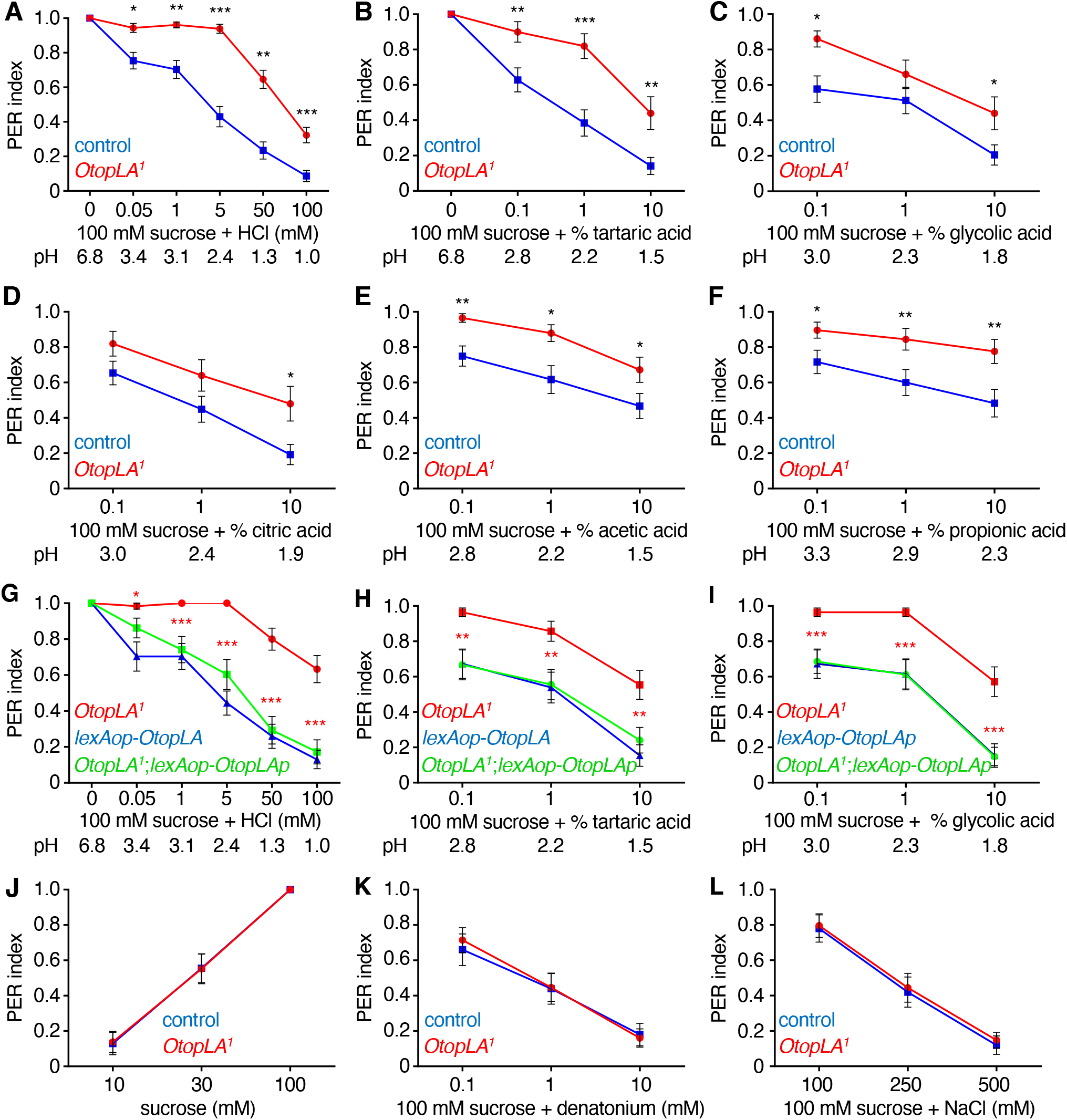
Mutation of *OtopLA* impairs gustatory repulsion to acids. (*A—F*) PER indexes obtained using control (*w*^*1118*^) and *OtopLA*^*1*^ flies upon stimulation of the labellum with 100 mM sucrose alone or 100 mM sucrose mixed with the indicated concentrations of acids. (*A*) Sucrose and HCl. Control, n=64. *OtopLA*^*1*^, n=65. (*B*) Sucrose and tartaric acid. Control, n=39. *OtopLA*^*1*^, n=25. (*C*) Sucrose and glycolic acid. Control, n=39. *OtopLA*^*1*^, n=25. (*D*) Sucrose and citric acid. Control, n=39. *OtopLA*^*1*^, n=25. (*E*) Sucrose and acetic acid. Control, n=30. *OtopLA*^*1*^, n=29. (*F*) Sucrose and proprionic acid. Control, n=30. *OtopLA*^*1*^, n=29. (*G—I*) Testing for rescue of the *OtopLa*^*1*^ defects in acid aversion by expressing the *OtopLAp* transgene under control of the *LexA* knocked into *OtopLA*^*1*^. The PERs were assayed from the following lines using the indicated concentrations of acids: 1) *OtopLA*^*1*^, 2) *lexAop-OtopLA*, and 3) *OtopLA*^*1*^;*lexAop-OtopLA* (rescue). Asterisks indicate significance differences between *OtopLA*^*1*^ and the rescue flies. (*G*) HCl. *OtopLA*^*1*^, n=30. *lexAop-OtopLA*, n=27. *OtopLA*^*1*^;*lexAop-OtopLA*, n=29. (*H*) Tartaric acid. *OtopLA*^*1*^, n=28. *lexAop-OtopLA*, n=26. *OtopLA*^*1*^;*lexAop-OtopLA*, n=28. (*I*) Glycolic acid. *OtopLA*^*1*^, n=28. *lexAop-OtopLA*, n=26. *OtopLA*^*1*^;*lexAop-OtopLA*, n=28. (*J*) Testing whether the *OtopLA*^*1*^ mutation impairs the PER response to sucrose. Control (*w*^*1118*^), n=27. *OtopLA*^*1*^, n=29. (*K*) Testing whether the *OtopLA*^*1*^ mutation impairs the PER response to denatonium mixed with 100 mM sucrose. Control (*w*^*1118*^), n=27. *OtopLA*^*1*^, n=29. (*L*) Testing whether the *OtopLA*^*1*^ mutation impairs the PER response to NaCl mixed with 100 mM sucrose. Control (*w*^*1118*^), n=27. *OtopLA*^*1*^, n=29. Mann-Whitney U tests. Error bars, s.e.m.s. *p < 0.05, **p < 0.01, and ***p < 0.001.

To test whether the *OtopLA*^*1*^ flies also show a deficit in rejecting sucrose with organic acids we conducted PER assays with tartaric acid, glycolic acid, citric acid, acetic acid and propionic acid. The degree of suppression of the PERs in control flies varies greatly with different carboxylic acids (Fig. 2 *B—F*) (8, 9, 15). Nevertheless, *OtopLA*^*1*^ flies exhibited decreased behavioral aversion towards each of the organic acids (Fig. 2 *B—F*).

The preceding analyses indicate that one or more of the five isoforms with the common translational start site (*OtopLAc-g*) functions in acid taste. Therefore, to identify the key isoform, we isolated proboscises, prepared mRNA, and performed RT-PCR using primers that amplify all the isoforms except *OtopLAa* (Fig. S2*A*). We isolated and sequenced eight cDNAs from the proboscis, all of which encoded a novel 948 amino acid protein (OtopLAp) that differed slightly from OtopLAc-g (Fig. S2*A*). OtopLAp is most similar to OtopLAg, except for missing residues 204 to 210 in OtopLAg, which are encoded by the small fifth coding exon of the *OtopLAg* mRNA, and the last 4 amino acids of the eighth exon (residues 909-912; Fig. S2*A*). To test if *OtopLAp* is sufficient to restore normal acid sensitivity to the *OtopLA*^*1*^ mutant, we generated flies expressing *lexAop-OtopLAp* and introduced the transgene into the *OtopLA*^*1*^ background so that it would be expressed under control of the *LexA* knocked into *OtopLA*^*1*^ (Fig. S2*A*). *OtopLAp* fully restored aversion to HCl, tartaric acid and glycolic acid (Fig. 2 *G—I*) demonstrating that *OtopLAp* is sufficient to confer acid taste.

### *OtopLA* mutants display a narrow behavioral deficit in tasting low pH

To evaluate whether *OtopLA* is specifically required for acid taste, we assayed the PERs to other chemicals. The attraction to sugar alone was identical between *OtopLA*^*1*^ and the control flies over a range of concentrations (Fig. 2*J*). Thus, the reductions in repulsion to sugars mixed with acids did not appear to be due to a change in sugar sensitivity. The aversion to the bitter compound, denatonium, and to high levels of NaCl exhibited by *OtopLA*^*1*^ flies were also indistinguishable from controls (Fig. 2 *K* and *L*). Thus, the *OtopLA*^*1*^ mutation did not cause a broad deficit in gustatory sensation.

### *OtopLA* mutants are impaired in acid-induced action potentials

If loss of *OtopLA* impairs acid sensitivity by disrupting reception in GRNs, then this should cause a decrease in acid-induced action potentials. The labellum is decorated with 31 gustatory hairs (sensilla) that fall into three length classes: small (S-type), intermediate (I-type) and large (L-type) (3). 9 out of the 11 S-type sensilla are categorized as either S-a or S-b sensilla on the basis of distinct sensitivities to bitter compounds (16). The two S-c-sensilla (S4 and S8) are unresponsive to all bitter chemicals tested (16). The S-b sensilla (S3, S5 and S9) have been reported to be responsive to acids (8). Using control flies, the electrolyte alone (30 mM tricholine citrate; pH 6.8) induced very few action potentials in S-b sensilla (Fig. 3 *A—C*). Consistent, with a previous study (8), HCl (pH 4 and pH 2) activated S-b sensilla (Fig. 3 *A—D* and *F*), but not S-a (e.g. S6 and S7; Fig. S3 *A* and *B*) or the S-c sensillum (e.g. S4; Fig. S3*C*). We observed little or no sensitivity to acids in the L sensilla (e.g. L7; Fig. S3*D*). Also as reported, the I-a type (e.g. I7) are not simulated by acid (Fig. S3*E*), while I-b (e.g. I8) type of I sensilla are responsive (8) (Fig. S4*F*).

To determine whether the *OtopLA*^*1*^ mutation caused a reduction in acid-induced action potentials we examined HCl at pH 4 and 2. We found that the frequencies of action potentials were diminished in all S-b sensilla, and to a greater extent in S3 and S9 (Fig. 3 *A—G*). We recorded from one of the I-b sensilla (I8) and found that the frequency of acid-induced action potentials was greatly reduced in *OtopLA*^*1*^ flies (Fig. S3*F*).

**Fig. 3.**
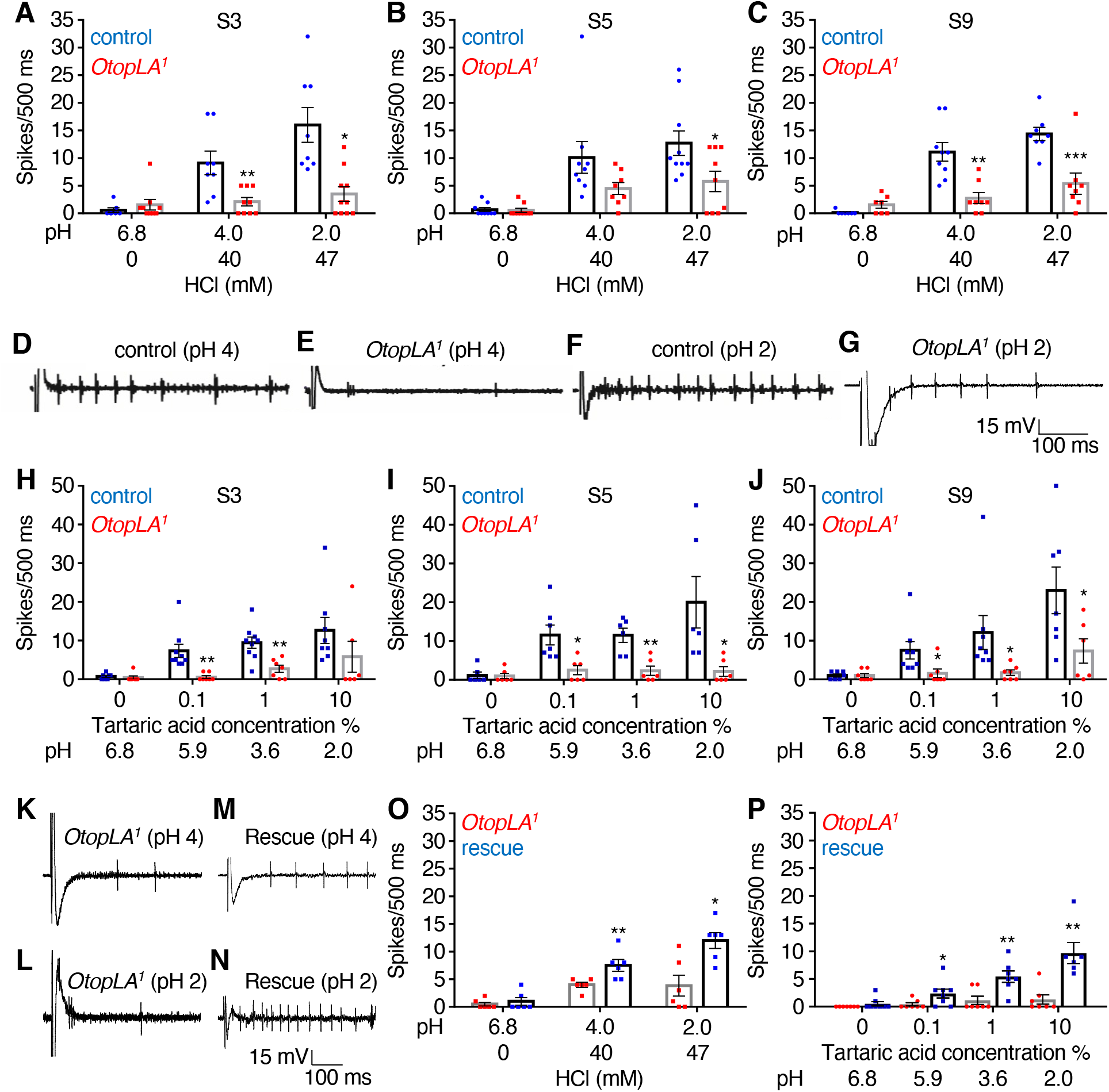
Tip recordings showing that mutation of *OtopLA* impairs acid-induced action potentials. (*A—C*) Tip recordings from the indicated s-b class sensilla (16) due to stimulation with HCl at the indicated pH values. Shown are the mean action potentials during the first 500 msec of the recordings with the control (*w*^*1118*^) and *OtopLA*^*1*^. The pH 6.8 solution contained only the electrolyte (30 mM 3 tricholine citrate). (*A*) Responses of S3 sensilla to HCl. Control, n=7—8. *OtopLA*^*1*^, n=9—10. (*B*) Responses of S5 sensilla to HCl. Control, n=9—10. *OtopLA*^*1*^, n=8—19. (*C*) Responses of S9 sensilla to HCl. Control, n=7—9. *OtopLA*^*1*^, n=7—8. (*D—J*) Representative traces for the first 500 msec obtained from S9 sensilla of control (*w*^*1118*^) and *OtopLA*^*1*^ flies during exposure to HCl at pH 4 or 2. (*H—J*) Tip recordings from the indicated s-b class sensilla due to stimulation with tartaric acid at the indicated pH values. Shown are the mean action potentials during the first 500 msec of the recordings with the control (*w*^*1118*^) and *OtopLA*^*1*^. (*H*) Responses of S3 sensilla to tartaric acid. Control, n=7—9. *OtopLA*^*1*^, n=6—7. (*I*) Responses of S9 sensilla to tartaric acid. Control, n=6—7. *OtopLA*^*1*^, n=6. (*J*) Responses of S3 sensilla to tartaric acid. Control, n=7—8. *OtopLA*^*1*^, n=6—7. (*K—N*) Testing for rescue of the deficit in HCl-induced action potentials in *OtopLA*^*1*^ flies expressing the *OtopLAp* transgene. Representative traces for the first 500 msec obtained from S9 sensilla of *OtopLA*^*1*^ and *OtopLA*^*1*^;*lexAop-OtopLAp* (rescue) flies upon stimulation with HCl at pH 4 or 2. (*O*) Assaying for changes in mean HCl-induced action potentials in *OtopLA*^*1*^ flies expressing the *OtopLAp* transgene. The mean action potentials were during the first 500 msec from S9 sensilla stimulation with HCl of the indicated pH values. *OtopLA*^*1*^, n=6. and *OtopLA*^*1*^;*lexAop-OtopLA* (rescue), n=6. (*P*) Assaying for changes in mean tartaric acid-induced action potentials in *OtopLA*^*1*^ flies expressing the *OtopLAp* transgene. The mean action potentials were during the first 500 msec from S9 sensilla stimulation with tartaric acid of the indicated pH values. *OtopLA*^*1*^, n=7. and *OtopLA*^*1*^;*lexAop-OtopLA* (rescue), n=6—8. Unpaired Student *t*-tests. Error bars, s.e.m.s. *p < 0.05, **p < 0.01, and ***p < 0.001.

We also compared the neuronal excitability of control and *OtopLA*^*1*^ over a range of concentrations of tartaric acid. The s-b and I-b taste sensilla from the mutant flies exhibited large reductions in in their responses to all levels of tartaric acid (Fig. 3 *H—J* and Fig. S3*G*). While *OtopLA* is required for normal acid-induced action potentials, it is difficult to directly correlate the effects of pH on behavior and action potentials since PERs involve integration of sugar-induced activation of A (sugar responsive) GRNs, and activation of a subset of B (bitter responsive) GRNs by acids (3). In addition, because the TCC electrolyte was used for the tip recordings, higher levels of acids were needed to achieve the same pH relative to when the acid is mixed with sucrose.

To test if the *OtopLAp* is sufficient to restore neuronal sensitivity to the *OtopLA*^*1*^ mutant, we conducted extracellular tip recordings. We created flies harboring the *lexAop-OtopLAp* transgene, and crossed it into the *OtopLA*^*1*^ background so that it was expressed under control of the *LexA*. We then assayed HCl-induced action potentials by recording from S9 sensilla. We found that introduction of the *OtopLAp* transgene significantly increased the frequency of HCl-driven action potentials in *OtopLA*^*1*^ flies (Fig. 3 *K—O*). Introduction of the *OtopLAp* transgene also increased the low sensitivity to tartaric acid in the S9 sensilla of the *OtopLA*^*1*^ mutant (Fig. 3*P*).

### *OtopLA* function is required in B GRNs for sensitivity towards acids

S-type sensilla house four types of GRNs including A GRNs that are activated by attractive tastants such as sugars, B GRNs that are stimulated by bitter compounds, acids and other aversive chemicals, C GRNs that respond to water, and D GRNs that are activated by cations (3). I-type sensilla harbor just A GRNs and B GRNs, while L-type sensilla house A, C, D and E (low salt) GRNs (3). Although B GRNs respond to acids (8), loss of *OtopLA* did not cause a general deficit in these GRNs since the frequency of denatonium-induced spikes were the same in control and mutant flies (Fig. S3 *H* and *I*).

Carboxylic acids have been reported to suppress feeding by activating B GRNs and suppressing the sugar response of A GRNs (8). We focused on citric acid and an L-type sensilla (L7), which houses A but not B GRNs, and found that citric acid suppressed sugar-induced action potentials in a concentration dependent manner as reported (Fig. S3*J*) (8). However, HCl did not suppress the sugar activation of A GRNs even at a pH 2.0 (Fig. S3*K*), which is lower than the pH of the highest concentration of citric acid tested (pH 2.2; Fig. S3*J*).

To identify the type of GRN that requires *OtopLA* for sensitivity towards acids, we performed gene silencing in different GRNs using the two *OtopLA* RNAi lines that we used to knock down *OtopLA* under the control of a pan-neuronal driver (*elav-GAL4*). We drove the *UAS-OtopLA-RNAi* transgenes in different classes of GRNs under control of cell-type specific *GAL4*s, and then conducted PER assays. Knockdown of *OtopLA* in A (*Gr5a-GAL4*), C (*ppk28-GAL4*), D (*ppk23-GAL4*) or E (*Ir94a-GAL4*) GRNs had no effect on the suppression of 100 mM sucrose appeal by 5 mM HCl (Fig. 4*A*). Taste hairs also contain an associated mechanosensory neuron (3). There was no effect resulting from RNAi knockdown of *OtopLA* in these neurons (*nompC-GAL4*). However, when we silenced *OtopLA* in B neurons (*Gr66a-GAL4*), the repulsion towards 5 mM HCl was suppressed greatly (Fig. 4*A*). To further characterize the effect of silencing *OtopLA* in B neurons, we conducted PER assays with a range of HCl concentrations (Fig. 4*B*). The outcome was the same as that obtained with pan-neuronal silencing of *OtopLA* –there was a general loss of sensitivity towards low pH across a range of concentrations (Fig. 4*B*).

**Fig. 4.**
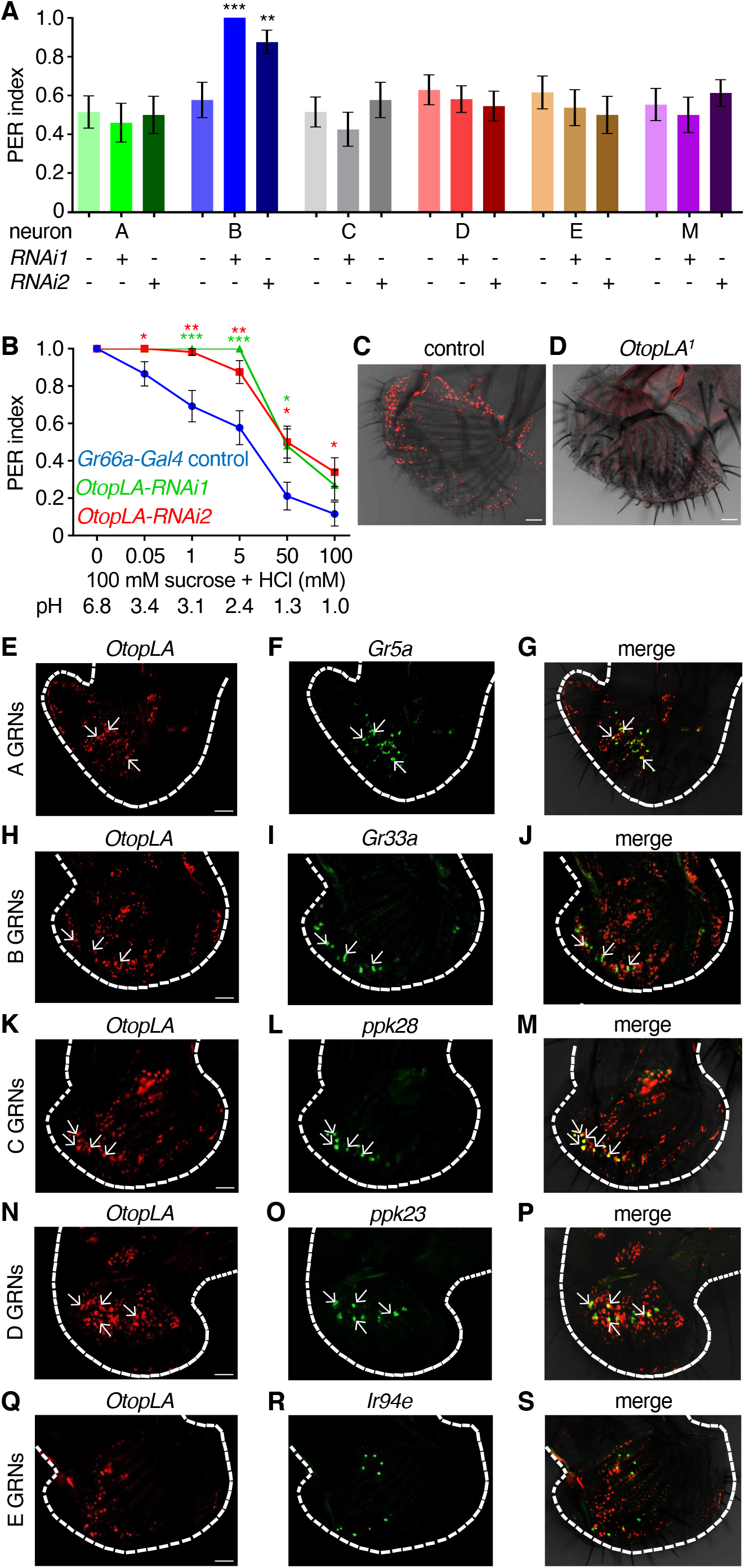
OtopLA functions in the B GRNs for detection of acids. (*A*) PER assays to test effects on acid aversion after RNAi knockdown of OtopLA in different classes of GRNs (A—E) and the mechanosensory neuron (M) associated with taste sensilla. PERs were in response to 100 mM sucrose mixed with 5 mM HCl. *OtopLA* was knocked down in A—E GRNs and M using two independent RNAi lines for each class. *GAL4* lines used to specifically express the RNAi lines in different neurons: *Gr5a-GAL4* for A GRNs, *Gr66a-GAL4* for B GRNs, *ppk28-GAL4* for C GRNs, *ppk23-GAL4* for D GRNs, *Ir94e-GAL4* for E GRNs and *nompC-GAL4* for M neurons. The controls without RNAi lines were the *GAL4* lines crossed to *w*^*1118*^. n≥25. (*B*) PER assays to test the effects of stimulation with 100 mM sucrose plus different concentrations of HCl due to RNAi knockdown of *OtopLA. Gr66a-GAL4* control, n= 26. *OtopLA-RNAi1*, n= 26. *OtopLA-RNAi2*, n= 28. (*C*) *In situ* hybridization of *OtopLA* transcripts in a control (*w*^*1118*^) labellum. Scale bar indicates 10 μm. (*D*) *In situ* hybridization of *OtopLA* transcripts in a *OtopLA*^*1*^ labellum. Scale bar indicates 10 μm. (*E—S*) Images of in situ hybridizations of *OtopLA* and the indicated reporters in labella from *w*^*1118*^ flies. In each row, the left panel indicates the distribution of *OtopLA* transcripts (red). The middle panel shows the markers for A—E GRNs (green). The right panel shows the merge of the *OtopLA* RNAs and marker RNAs. Scale bars indicate 10 μm. Mann-Whitney U tests. Error bars, s.e.m.s. *p < 0.05, **p < 0.01, and ***p < 0.001.

To examine expression in the labellum we used two approaches. We first took advantage of the *LexA* reporter knocked into the *OtopLA*^*1*^ allele (Fig. S2*A*) to drive expression of *lexAop-mCherry*. Although the mCherry signal was modest, it appeared to label GRNs associated with both taste pegs (Fig. S4*A*), which are flat sensilla situated between the pseudotrachea (17), and taste hairs (Fig. S4*D*). However, when we used the *poxneuro*^*70*^ (*poxn*^*70*^) mutation to eliminate all chemosensory hairs without effecting peg GRNs (18) we found that neither 5 mM HCl nor 1% tartaric acid was effective in suppressing the attraction to sucrose (Fig. S4*B*). Conversely, when we inactivated peg GRNs only (*57F03-GAL4*and *UAS-kir2*.*1*) (19), there was normal acid induced suppression of the sugar response (Fig. S4*C*). Thus, the peg GRNs did not appear to be required for the acid response.

Due to the weak reporter signals, in order to determine which type of neurons express *OtopLA*, we performed *in situ* hybridizations. We first probed the labellum with the *OtopLA* probe alone and observed RNA signals in control but not *OtopLA*^*1*^ labella (Fig. 4 *C* and *D*). However, the labeling of any given labellum was not complete since the signals were limited by penetration of the probe. Next, we performed double *in situ* hybridization experiments using markers for the five different GRN subsets (A—E; Table 1), which have different response profiles to tastants (3). ∼50% of the *OtopLA* positive neurons overlapped with the A—D GRNs, and none overlapped with E GRNs (Fig. 4 *E—S*; Table 1). The remaining neurons may be peg GRNs. However, as described above, expression of *OtopLA* in peg GRNs does not contribute to acid taste. Since *OtopLA* functions in B GRNs, to further examine the overlap of *OtopLA* with B GRNs, we used the *LexA* knocked into *OtopLA*^*1*^ to drive expression of *lexAop-GFP*. When we labeled the B GRNs with the *Gr66a-GAL4*, which drove expression of *UAS-td-tomato*, we found that a subset of the GFP-positive GRNs (*OtopLA*) were also labeled by td-Tomato, although the percentage (∼12%) was higher than obtained using *in situ* hybridizations (Fig. S4 *D—F*).

**Table 1.** GRNs expressing *OtopLA* and markers for A—E classes of GRNs (3). Double *in situ* hybridizations were performed using RNAscope (ACD, Hayward, CA, USA). Mean number of GRNs in proboscises (n≥3) expressing *OtopLA* and the indicated A—E markers. The average number of GRNs expressing *OtopLA* varied in different double-labeling experiments due to variations in probe penetration.

| GRN | Former name | GRN maker | GRNs labeled by marker | GRNs labeled by *OtopLA* | *OtopLA<sup>+</sup>* GRNs labeled by marker | % *OtopLA<sup>+</sup>* GRNs labeled by marker |
| --- | --- | --- | --- | --- | --- | --- |
| A | sweet | *Gr5a* | 24.0 ± 1.6 | 75.6 ± 3.3 | 5.6 ± 0.3 | 7.4 |
| B | bitter | *Gr33a* | 18.6 ± 1.3 | 83.0 ± 3.5 | 5.0 ± 0.5 | 6.0 |
| C | water | *ppk28* | 16.0 ± 0.6 | 66.0 ± 3.0 | 11.6 ± 0.3 | 17.5 |
| D | cation | *ppk23* | 16.6 ± 1.2 | 71.6 ± 4.3 | 14.2 ± 0.6 | 19.7 |
| E | low salt | *Ir94a* | 9.3 ± 0.6 | 70.0 ± 4.6 | 0.0 | 0 |

Since expression of *OtopLA* was observed in neurons required for tasting appetitive tastants, we investigated the possibility that *OtopLA* is required for the attraction to low concentrations of acids. However, we did not observe significant attraction to low concentrations of HCl (Fig. S4*G*), and the PER response to low concentrations of HCl were similar in control and *OtopLA*^*1*^ flies (Fig. S4*G*).

## Discussion

The functional conservation of the Otop channels for acid taste in flies is striking given that chemosensory receptors tend to vary greatly in flies and mammals (4), which diverged ∼800 million years ago. In contrast to the Otop channels, the two major families of fly receptors (GRs and IRs), which function in tasting sugars, bitter compounds, acetic acid, amino acids, polyamines, DEET, CO2 and other tastants are not present in mammals (3, 20). The retention of Otop channels for acid taste in flies and mice is remarkable since the gross anatomies of the gustatory systems are very different (4). In addition, the taste receptor cells in flies are neurons, while they are modified epithelial cells in mammals (4).

The common role for Otop proteins for acid taste in flies and mammals cannot be explained by greater selective pressure for maintaining a receptor for a mineral (e.g. H^+^) versus organic molecules since other minerals (Ca^2+^ and Na^+^) are sensed in flies through IRs, which are not present in mammals (21-23). Thus, the retention of Otop channels for acid taste in flies and mammals underscores the very strong selection for this acid sensor for animal survival. Otop-related proteins are encoded in many distantly related terrestrial and aquatic vertebrates ranging from the platypus to frogs, and pufferfish, as well as ancient invertebrates such as worms and insect disease vectors, including *Aedes aegypti* (12, 24). Thus, despite the considerable diversity of most chemosensory receptors, it is plausible that Otop channels endow a large proportion of the animal kingdom with acid taste.

A question concerns the cellular mechanism through which the sensation of protons is encoded. It has been reported that acids cause repulsion of sugary foods by activation of B GRNs and suppression of A GRNs by sugars (8). This previous study focused on carboxylic acids and we repeated this finding with citric acid. However, when we decreased the pH of sucrose with HCl, we did not observe reduced sucrose-induced action potentials. Thus, we conclude that protons do not suppress A GRNs. Rather we suggest that A GRNs are inhibited by certain organic anion moieties of carboxylic acids. A mechanism by which the activities of both A GRNs and B GRNs are affected by carboxylic acids, but only B GRNs by protons could provide a coding mechanism for differentiating between protons and carboxylic acids. Consistent with the role of B GRNs but not A GRNs in responding to HCl, we found that OtopLA, functions specifically in B GRNs. Since low levels of certain carboxylic acids are attractive, it is intriguing to speculate that there exists another channel in A GRNs that contributes to acid attraction. Another future issue concerns the mechanism through proton influx through OtopLA induces action potentials in B GRNs. In mammalian taste receptor cells, protons inhibit a K^+^ channel (K_IR_2.1), which contributes to depolarization (25). It is intriguing to speculate that a similar mechanism might contribute to depolarization and activation of the voltage-gated Na^+^ channels in *Drosophila*.

## Materials and Methods

### *Drosophila* stocks

Flies were reared at 22°C to 25°C in standard media. Control flies were *w*^*1118*^ unless otherwise mentioned. The following lines were obtained from the *Drosophila* Bloomington Stock Center: *w*^*1118*^ (BL 5905), *nompC-GAL4* (BL 50131), *Ir94e-GAL4* (BL 60725), *elav-GAL4;UAS-Dcr2* (BL 25750), *Gr5a-GAL4* (BL 57592), *lexAop-GFP* (BL 32207), *lexAop-RFP* (BL67093),*UAS-tdTomato* (BL 36328), *Poxn*^*70*^ (BL 60688), *UAS-Kir2*.*1* (BL 6596) and *57F03-GAL4* (BL 46386). The following lines were obtained from the Vienna Drosophila *RNAi* Center: *OtopLA-RNAi1* (v104973), *OtopLA-RNAi2* (v100847), *OtopLB-RNAi1* (v101936), *OtopLB-RNAi2* (v3452), *OtopLC-RNAi1* (v47248) and *OtopLC-RNAi2* (v19613). The following lines were obtained were obtained from other investigators: *ppk23-Gal4* (26), *ppk28-GAL4* (27), and *Gr66a-GAL4* (28).

### Chemicals

Sucrose (84097), denatonium (D5765), tartaric acid (251380), glycolic acid (124737), propionic acid (81910), acetic acid (A6283) and citric acid (C2404) were obtained from Sigma. NaCl (S271-3) and HCl (A144-212) were obtained from Fisher Scientific. For PER assays, water was used as the solvent. Tastants were dissolved in 30 mM TCC (Sigma, T0252) for extracellular tip recordings.

### Generation of *OtopLA*^*1*^ and *OtopLA*^*2*^ mutant flies

To create the *OtopLA*^*1*^ and *OtopLA*^*2*^ mutants we used CRISPR/Cas9. To generate *OtopLA*^*1*^, we replaced 40 base pairs at the beginning of the coding sequence of *OtopLA* with the *LexA* and *mini-white* (*w*^*+*^) genes (Fig. S2*A*). The deletion removed first 14 amino acids and also changed the reading frame. The guide RNAs for creating this line were: guide RNA1: gcgaggggaaaggaatgcagcgg and guide RNA2: tgagatgcgcgaaagattactgg. The PCR primers used to generate the 5’ and 3’ homology arms were: forward primer for 5’ arm: gacgcataccaaacggtaccaatgctcccgatttgctggctagc, reverse primer for 5’ arm: ttttgattgctagcggtacctcctttcccctcgctcagc, forward primer for 3’ arm: ctaggcgcgcccatatgtactggaccagccgcggg, and reverse primer for 3’ arm: gacaagccgaacatatgggtgggcaggactcgtg

To generate *OtopLA*^*2*^, we replaced the first 52 base pairs at the beginning of the coding sequence of *OtopLAa* with the *LexA* and *mini-white* (*w*^*+*^) genes (Fig. S2*A*). The deletion removed first 18 amino acids and changed the reading frame. The guide RNAs for creating this line were: guide RNA1: acggaaacggaaacaatgggcgg and guide RNA2: cgtcgagggcggggacaatatgg. The PCR primers used to generate the 5’ and 3’ homology arms were: forward primer for 5’ arm: gacgcataccaaacggtacccagggcccgtttcagtt, reverse primer for 5’ arm: ttttgattgctagcggtacctgtttccgtttccgtttcggtct, forward primer for 3’ arm: ctaggcgcgcccatatgatatggccaccttgccgg, and reverse primer for 3’ arm: gacaagccgaacatatgaaccccagacagtgtgcag

The plasmids to create *OtopLA*^*1*^ and *OtopLA*^*2*^ were injected by Bestgene into the *vas-cas9* background (BL 51324). After confirming the mutations by PCR (forward primer: agtcgttgccaggatagacg reverse primer: catcgtggggaactcatcat) and DNA sequencing, we outcrossed both mutant lines to the *w*^*1118*^ background for 6 generations.

### Generation of *lexAop-OtopLAp* transgenic flies

50 *w*^*1118*^ proboscises were dissected under liquid N2 and RNA was extracted using the Qiagen RNeasy mini kit (74104). cDNAs were synthesized from freshly extracted mRNA using Superscript IV Mastermix with the ezDNase enzyme (Invitrogen 11766050). The entire coding sequence of the *OtopLAp* isoform was PCR amplified (34 cycles at an annealing temperature 56°C) from freshly synthesized cDNAs using the following primers: forward primer: gcggccgcggctcgagaaaggaatgcagcggtgt, and reverse primer: acaaagatcctctagattactccagacgt. The *OtopLAp* cDNA was cloned between the XhoI and XbaI sites of the *lexAop* vector (Addgene 26224) (29). The plasmid was injected by Bestgene into the *attP2* background (BL 8622), which has an *attP* docking site (68A4) on the 3^rd^ chromosome to create *lexAop-OtopLA* transgenic flies. The transgenic line was verified by PCR.

### Proboscis Extension Response (PER) assays

To perform PER assays (30), we collected flies that were 0-2 days old, and aged them in vials containing 10 males and 10 females until they were 5-7 days old. The flies were starved on water-soaked Kimwipes for 24-26 hours prior to the experiments. The flies were fitted within P-200 tips truncated in a manner to allow only the head to protrude outside. The other end of the tip was sealed with clay. Paper wicks were used to present the taste stimuli to the flies. We made contact for ∼2 seconds and scored the fly’s response over the following 7 seconds. The flies were satiated with water at the beginning of the experiment and between consecutive stimuli. In addition, the flies were tested with 100 mM sucrose both at the beginning and the end of the experiments. Only those flies that gave a positive response in both cases were considered. After every positive response, flies were tested with water to determine whether the PER was due solely to the tastants. Flies that continued to drink water for ≥1 minute were discarded. Complete proboscis extensions were scored as 1 while partial extensions were scored as 0.5. A score of 0 was recorded on no extension. The mean score obtained from all individual flies tested was calculated as PER index. PER experiments were done blinded except in cases where there were differences in eye colors between the lines that precluded us from doing so.

### Extracellular tip recordings

Extracellular tip recordings were conducted as previously described (31). 7-10 day-old flies were used for the recordings. The tastants were dissolved in 30 mM tricholine citrate (TCC), which served as the electrolyte. The taste solutions were back-filled into recording electrodes (World Precision Instruments, 1B150F-3). Action potentials induced by the tastants were amplified and digitalized using an IDAC-4 data acquisition software. Autospike software (Syntech) was used to visualize and manually count the spikes. Neuronal responses were quantified by counting the number of spikes in the first 500ms following contact with the stimulus.

### Immunostaining

Proboscises were dissected in 1% PBS and fixed in 4% paraformaldehyde for 30 minutes. They were then washed with PBST (PBS+0.3% Triton X-100) for 1 hour (3 washes of 20 minutes each), and blocked using normal goat serum for 1 hour. The whole mount tissues were stained in primary antibodies in blocking buffer for 3-4 days, washed with PBST for one hour and subsequently stained using secondary antibodies in blocking buffer for 2-3 days. The tissues were mounted on glass slides with Vectashield mounting media and images were acquired using a Zeiss LSM 700 confocal microscope. Primary antibodies: anti-GFP (chick 1:20) and anti-DsRed (rabbit, 1:50, Takara Bio#632496). Secondary antibodies: Alexa Fluor 488 conjugated goat anti-chicken (Thermo Fisher Scientific, A-11039) and Alexa Fluor 568 conjugated goat anti-rabbit (Thermo Fisher Scientific, A-11036).

### mRNA hybridizations

To localize mRNAs in proboscis whole-mount tissue, we adapted RNAscope using primers to *OtopLA, Gr5a, Gr33a, ppk28, ppk23* and *Ir94e*, which were designed by Advanced Cell Diagnostics (ACD, Hayward, CA, USA). ∼10 proboscis were dissected in PBS, placed in 1.5 mL Eppendorf tubes, washed once with PBS and fixed in 4% paraformaldehyde in 1 mL PBS for 16 hour at 4°C. The tissue was then immersed in a series of 10%, 20% and 30% sucrose, each time allowing the tissue to sink the bottom of the tube. Then the tissue was washed in PBS, re-fixed in 4% paraformaldehyde in PBS at room temperature for 10 minutes and washed for 3×5 minutes in PBS. The samples were then dehydrated in a series of 50%, 75% and 100% ethanol in PBS. The ethanol was removed completely, and the tissue was air-dried at room temperature for 30 minutes. The tissue was treated with 3% hydrogen peroxide in PBS for 10 min to inactivate endogenous peroxidase activity and incubated in 50μL RNAscope Protease III for 30 min. Hybridizations with 100 μL each of the *OtopLA* and marker probes were performed overnight at 40°C in ACD HybEZ^™^ Hybridization System (110VAC) (ACD cat. no. 321461). The tissue was washed in an RNAscope wash buffer (ACD cat. no. 310091) for 3 × 2 minutes. The tissue was then incubated in a series of amplifier solutions (Amp’s) provided in the RNAscope® Multiplex fluorescent V2 assay kit (ACD cat. no. 323100, Newark, CA, USA) according to the manufacturer’s instructions. The tissue was incubated in 100 μL Amp1 for 2 hours at 40°C, in Amp2 for 2 hours at 40°C, Amp3 for 1 hour at 40°C, and C1 for 2 hours at 40°C. Between each step the tissue was washed for 5×3 minutes with wash buffer at room temperature. For fluorescent labeling, a working Opal dye solution was made fresh using a 1:500 ratio of Opal dye (Akoya Biosciences, Marlborough, MA, USA) to TSA buffer. 150 μl of working solution was added to each tube containing 1∼0 proboscises and incubated at 40°C for 2 hours, washed one in a wash buffer and mounted in Vectashield.

### Quantification and statistical analysis

Descriptions, results and sample sizes of each test are provided in the Figure legends. All replicates were biological replicates using different flies. Data for all quantitative experiments were collected on at least three different days. For the PER behavioral experiments each ‘‘n’’ represents an individual fly. Based on our experience and common practices in this field, we used a sample size of n ≥ 25 trials for each genotype or treatment for the PER assays. Each ‘‘n’’ for the tip recording experiments represents an analysis of a single, independent fly (n > 6). GraphPad Prism 8 software or MS Excel was used for statistical tests. We used unpaired Student’s *t*-tests for parametric tests (in MS Excel) and Mann-Whitney U test for non-parametric tests. Sample sizes were determined based on previous publications and are cited in the Figure legends. In all graphs, error bars indicate the standard error of the mean (s.e.m.). We set the significance level, a = 0.05. Asterisks indicate statistical significance: *p < 0.05, **p < 0.01, and ***p < 0.001.

## Author contributions

A.G. A.C, H.T and S.W. conducted the experiments, A.G., E.L and C.M. designed the experiments, and A.G. and C.M. wrote the initial draft of the paper, and all authors edited the manuscript.

## Acknowledgment

This work was supported by a grant to C.M. from the National Institute on Deafness and other Communication Disorders (DC007864).

## Competing interests

The authors declare no competing interest.

## Supplemental Figure legends

**Fig. S1.**
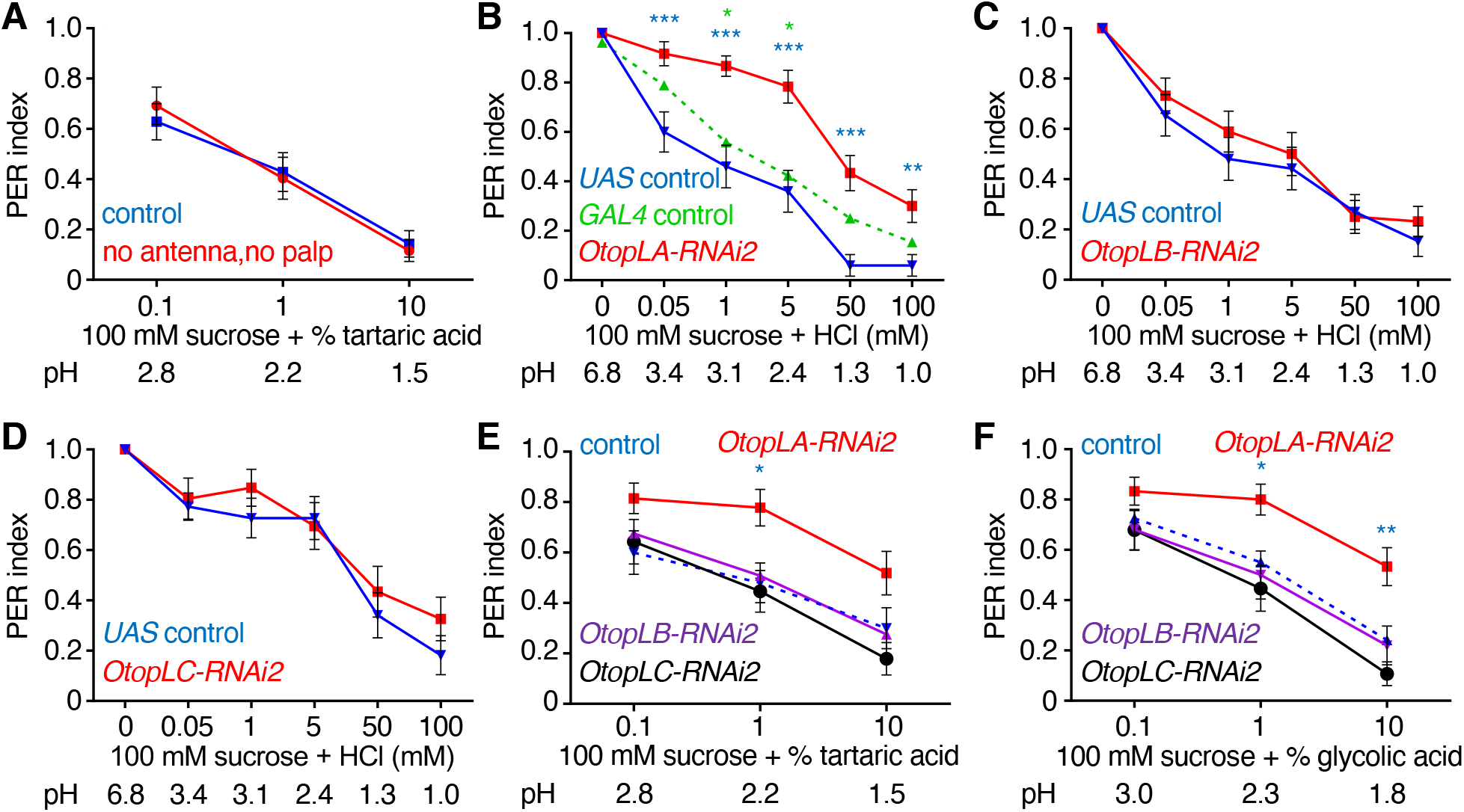
Screen with second set of RNAi lines further supports that *OtopLA* is required for aversion of acid taste. PER assays using either 100 mM sucrose alone or sucrose plus the indicated concentrations of HCl (mM) or organic acids (%). All RNAi lines (B— F) were generated by crossing the indicated *UAS-RNAi* lines to *elav-GAL4;UAS-Dcr2* flies. (*A*) Effects of removing olfactory organs on the PER responses to the indicated concentrations of tartaric acid. *w*^*1118*^ flies (intact flies) and *w*^*1118*^ flies in which the antenna and maxillary palp were surgically removed (no antenna, no palp). Intact flies, n=35. No antenna and no maxillary palp, n=26. (*B—F*) Effects of RNAi knockdown of different *OtopL* genes on the PER responses to the indicated acids. (*B*) Effect of knockdown with *OtopLA*-RNA2 on the PER to sucrose plus HCl. *OtopLA-RNAi2* is *UAS-OtopLA-RNAi2* (v100847) crossed to *elav-GAL4;UAS-Dcr2* flies (n=32). The *UAS* control (n=26) and *GAL4* control (n=25) are generating by crossing *w*^*1118*^ to either *UAS-OtopLA-RNAi2* or *elav-GAL4;UAS-Dcr2*, respectively. The blue and green asterisks indicate statistically significance differences between *OtopLA* silenced flies and the *UAS* and *GAL4* controls, respectively. The results with the *GAL4* control (green dotted line) are the same as that used in Fig. 1*B*. (*C*) Effect of knockdown with *OtopLB-RNAi2* on the PER to sucrose plus HCl. *OtopLB-RNAi2* is *UAS-OtopLB-RNAi2* (v3452) crossed to *elav-GAL4;UAS-Dcr2* flies (n=27). *UAS* control (n=26). (*D*) Effect of knockdown of *OtopLC-RNAi2* on the PER to sucrose plus HCl. *OtopLC-RNAi1* is *UAS-OtopLC-RNAi* (v19613) crossed to *elav-GAL4;UAS-Dcr2* flies (n=26). *UAS* control (n=27). (*E*) Effect of knockdown with *OtopLA-RNAi2* (n=30), *OtopLB-RNAi2* (n=25) and *OtopLC-RNAi2* (n=28) on PERs using the indicated concentrations of tartaric acid. *OtopLA-RNAi1* (n=30), *OtopLB-RNAi1* (n=25), *OtopLC-RNAi1* (n=28). The *GAL4* “control” is *elav-GAL4;UAS-Dcr2* flies (n= 65) and is also presented in Fig. 1*E*. The blue asterisks indicates a significant differences from the control. (*F*) Effect of knockdown with *OtopLA-RNAi2* (n=30), OtopLB-RNAi2 (n=25) and *OtopLC-RNAi2* (n=28) on PERs using the indicated concentrations of glycolic acid. The *GAL4* “control” is *elav-GAL4;UAS-Dcr2* flies (n= 65) and is also presented in Fig. 1*E*. Mann-Whitney U tests. Error bars, s.e.m.s. *p < 0.05, **p < 0.01, and ***p < 0.001.

**Fig. S2.**
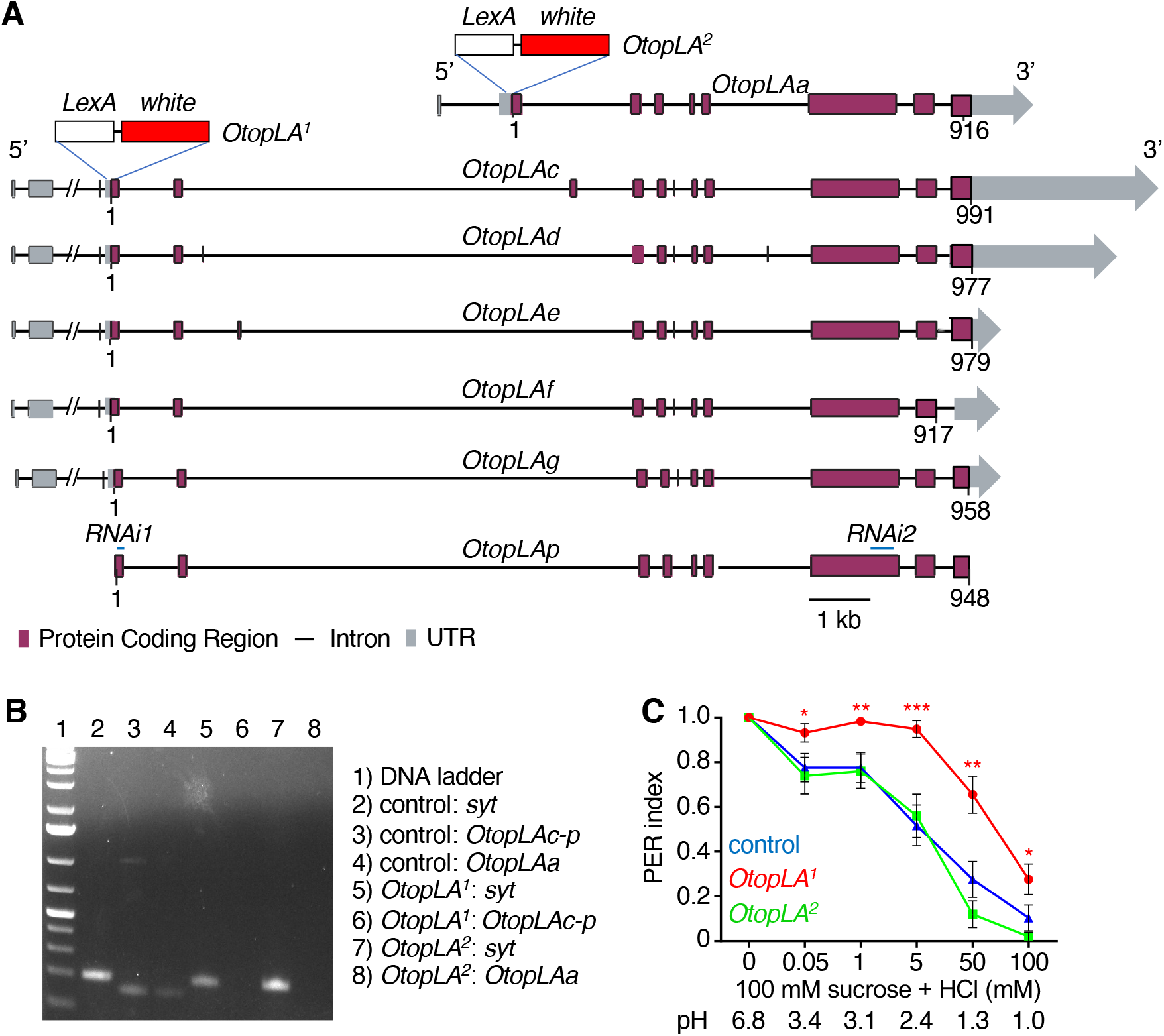
*OtopLA* mRNA isoforms and structures of *OtopLA*^*1*^ and *OtopLA*^*2*^ mutants. (*A*) Schematic of the *OtopLA* isoforms and the for generation of *OtopLA*^*1*^ and *OtopLA*^*2*^ mutants. The protein coding exons are indicated in purple and the horizontal lines indicate introns. To create *OtopLA*^*1*^ the first 40 base pairs of the coding sequence common to *OtopLAc*—*OtopLAg* and *OtopLAp* were replaced with *lexA* and *mini-white*. To generate *OtopLA*^*2*^, the first 52 base pairs of the coding region of *OtopLAa* were replaced with *lexA* and *mini-white*. The regions targeted by mRNA targeted by *RNAi1* and *RNAi2* are indicated. (*B*) RT-PCR using cDNAs synthesized from heads of control, *OtopLA*^*1*^ and *OtopLA*^*2*^. Lane 1: GeneRuler 1 kb Plus DNA Ladder (Thermo Fisher, SM1331). Lanes 2, 5 and 7 show *syt1* RT-PCR products (192 bp) from control, *OtopLA*^*1*^ and *OtopLA*^*2*^ cDNAs (primers: forward, tctggtcgtgcttcgagaag; reverse, cggatccctatgtcaaggtg). Lanes 3 and 6 are *OtopLAc-p* RT-PCR products using control (143 bp band) and *OtopLA*^*1*^ (no band) cDNAs, respectively (primers: forward, cagcggtgtccctacatt; reverse, ccatcgccctgcagctggttg). Lanes 4 and 8 are *OtopLAa* RT-PCR products from control (141 bp) and *OtopLA*^*2*^ (no band), respectively (primers: forward, atgggcggcggtgaagtgaaggt; reverse, ttccatctccttgttggcgg). (*C*) PER assay showing that *OtopLA*^*2*^, which specifically disrupts OtopLAa, does not impact on the aversion to HCl. Control (*w*^*1118*^), n=29. *OtopLA*^*1*^, n=29. *OtopLA*^*2*^, n=25. Mann-Whitney U tests. Error bars, s.e.m.s. *p < 0.05, **p < 0.01, and ***p < 0.001.

**Fig. S3.**
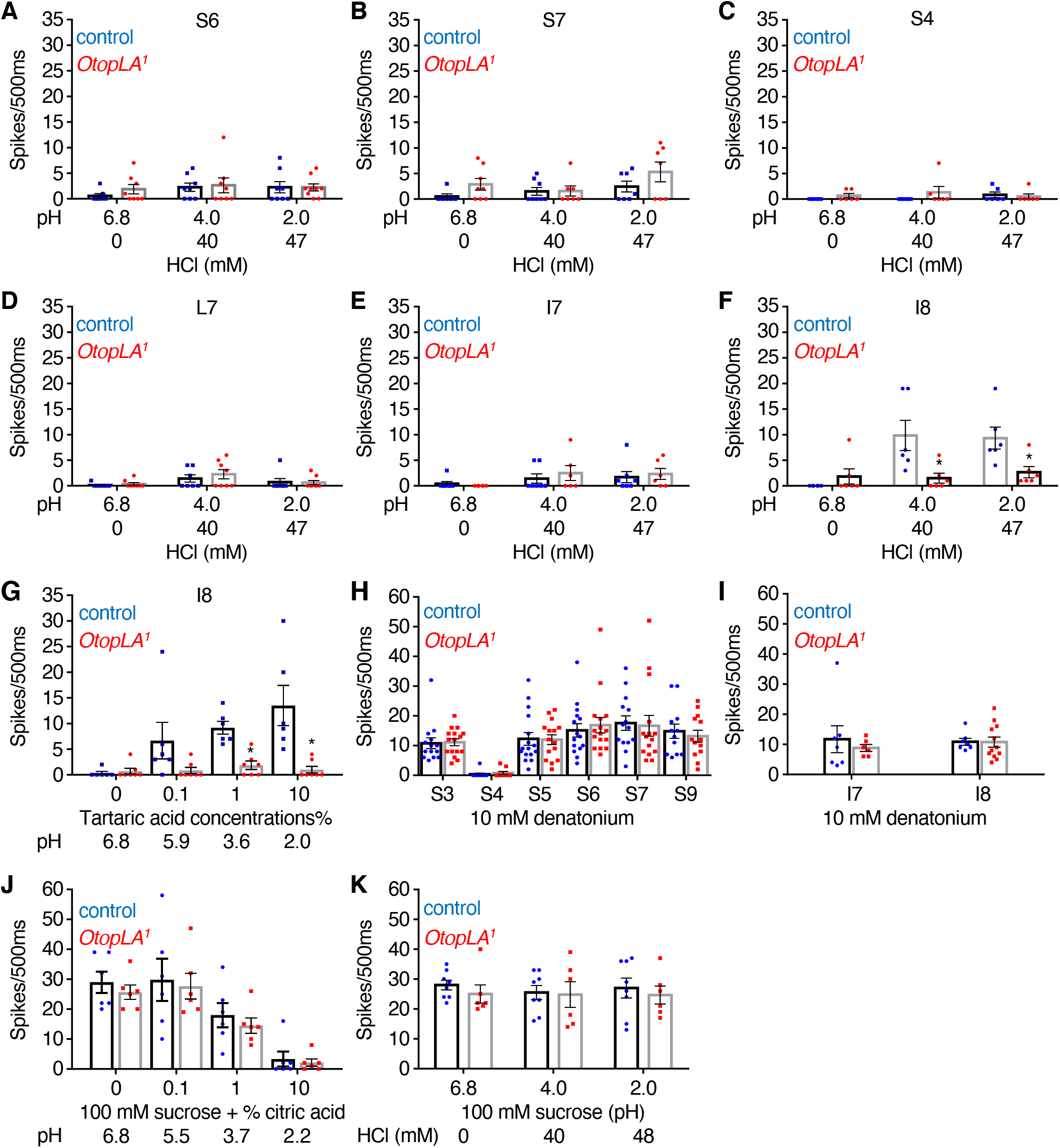
Tip recordings to test control and *OtopLA*^*1*^ responses of sensilla to acids, denatonium and sucrose. Mean action potentials elicited by the control (*w*^*1118*^) and the *OtopLA*^*1*^ mutant during the first 500 msec upon stimulating the indicated sensilla with the indicated chemical. (*A—F*) Stimulation of the indicated sensilla with HCl solutions at the indicated pH. The pH 6.8 solution contained only the electrolyte (30 mM 3 tricholine citrate). (*A*) Responses of S6 sensilla to HCl. Control, n=7—8. *OtopLA*^*1*^, n=8—9. (*B*) Responses of S7 sensilla to HCl. Control, n=6—8. *OtopLA*^*1*^, n=6—8. (*C*) Responses of S4 sensilla to HCl. Control, n=6. *OtopLA*^*1*^, n=6. (*D*) Responses of L7 sensilla to HCl. Control, n=7. *OtopLA*^*1*^, n=8. (*E*) Responses of I7 sensilla to HCl. Control, n=7. *OtopLA*^*1*^, n=6. (*F*) Responses of I8 sensilla to HCl. Control, n=6. *OtopLA*^*1*^, n=6. (*G*) Responses of I8 sensilla to the indicated concentrations of tartaric acid. Control, n=6. *OtopLA*^*1*^, n=6. (*H*) Responses of indicated S-type sensilla to 10 mM denatonium. Control, n≥12. *OtopLA*^*1*^, n≥12. (*I*) Responses of indicated I-type sensilla to 10 mM denatonium. Control, n≥6. *OtopLA*^*1*^, n≥6. (*J*) Responses of L7 sensilla to 100 mM sucrose mixed with citric acid of the indicated pHs. Control, n=6. *OtopLA*^*1*^, n=6. (*K*) Responses of L7 sensilla to 100 mM sucrose plus HCl at the indicated pHs. Control, n=8. *OtopLA*^*1*^, n=6. Unpaired Student’s *t*-tests. Error bars, s.e.m.s. *p < 0.05, **p < 0.01, and ***p < 0.001.

**Fig. S4.**
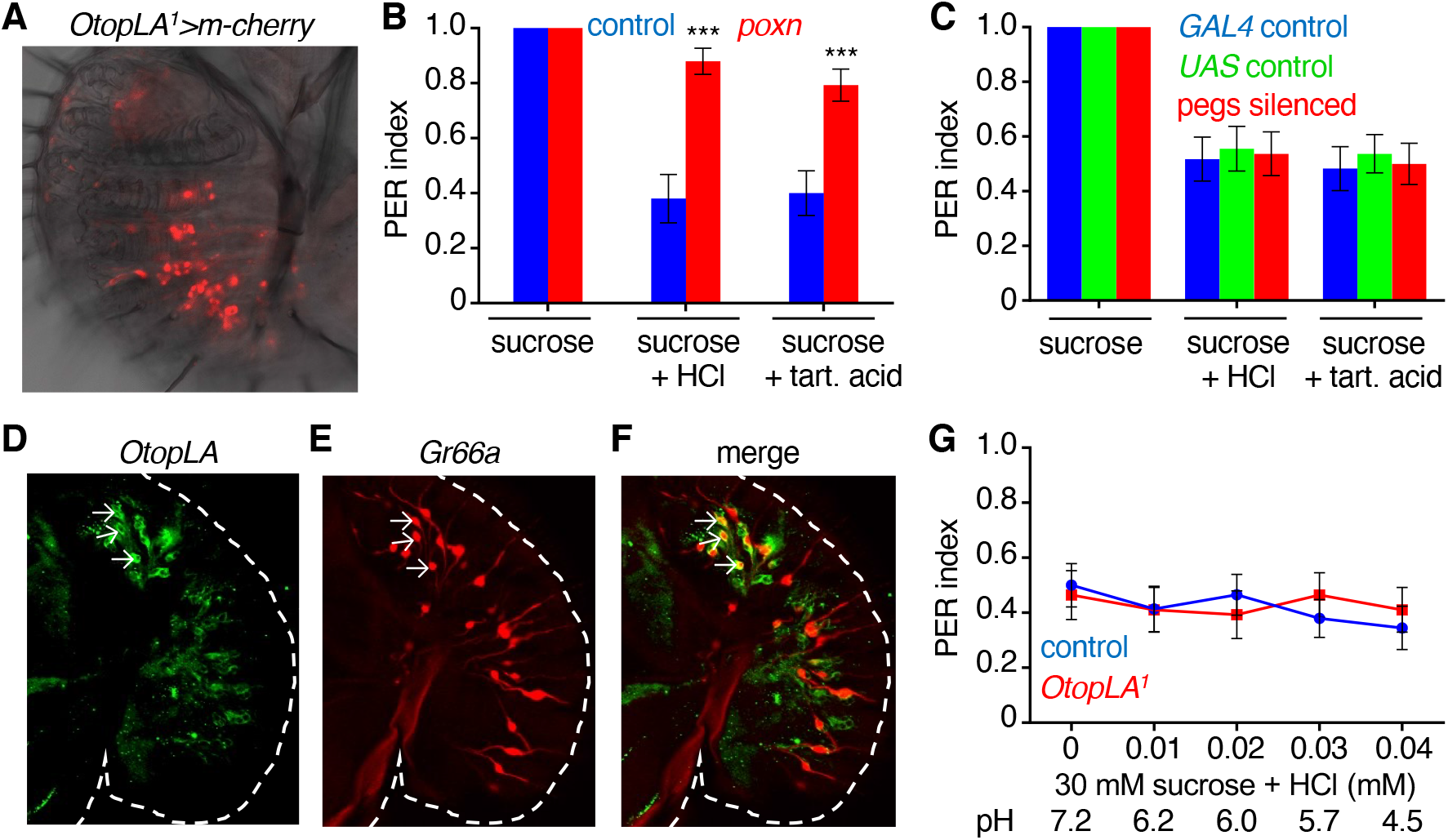
*OtopLA-LexA* reporter, peg GRNs are not required for acid taste, and low HCl levels do not affect the attraction to sucrose. (*A*) Confocal image of *OtopLA*^*1*^ (*lexA*) neurons (red; anti-DsRed) in the proboscis. The image focuses on neurons in taste pegs. Genotype: *OtopLA*^*1*^;*UAS-mCherry*. (*B*) PERs elicited by control (*w*^*1118*^) and *poxn*^*70*^ flies in response to stimulation with 100 mM sucrose, 100 mM sucrose mixed with either with 5 mM HCl or with 1% tartaric acid. Control, n=25. *poxn*^*70*^, n=29. (*C*) PER elicited from flies in the which the peg neurons are silenced show normal aversion to acids. Labella were stimulated with 1) 100 mM sucrose, 2) 100 mM sucrose mixed with 5 mM HCl, or 3) 100 mM sucrose plus 1% tartaric acid. *GAL4* control: *57F03-GAL4/+*, n=29. *UAS* control: *UAS-Kir2*.*1/+*, n=27. Pegs silenced: *UAS-Kir2*.*1/+;57F03-GAL4/+*, n=27. (*D—F*) Confocal image testing for overlap between the *OtopLA-LexA* and *Gr66a* reporters. The images focused on neurons in taste hairs. (*D*) *OtopLA*^*1*^ (*lexA*) was visualized using anti-GFP (green). Genotype: *OtopLA*^*1*^(*lexA*);*Gr66a-GAL4;UAS-tdTomato*/*lexAop-GFP*. (*E*) *Gr66a-GAL4* visualized using anti-DsRed (red). Genotype: *OtopLA*^*1*^(*lexA*);*Gr66a-GAL4;UAS-tdTomato*/*lexAop-GFP*. (F) Merge of *D* and *E*. (*G*) PER assays using 30 mM sucrose and low concentrations of HCl. Control (*w*^*1118*^), n=29. *OtopLA*^*1*^, n=28. Mann-Whitney U tests. Error bars, s.e.m.s. *p < 0.05, **p < 0.01, and ***p < 0.001.

## Notes

### Competing Interest Statement

The authors have declared no competing interest.

